# Identification of DNA motifs that regulate DNA methylation

**DOI:** 10.1101/573352

**Authors:** Mengchi Wang, Kai Zhang, Vu Ngo, Chengyu Liu, Shicai Fan, John W Whitaker, Yue Chen, Rizi Ai, Zhao Chen, Jun Wang, Lina Zheng, Wei Wang

## Abstract

DNA methylation is an important epigenetic mark but how its locus-specificity is decided in relation to DNA sequence is not fully understood. Here, we have analyzed 34 diverse whole-genome bisulfite sequencing datasets in human and identified 313 motifs, including 92 and 221 associated with methylation (methylation motifs, MMs) and unmethylation (unmethylation motifs, UMs), respectively. The functionality of these motifs is supported by multiple lines of evidences. First, the methylation levels at the MM and UM motifs are respectively higher and lower than the genomic background. Second, these motifs are enriched at the binding sites of methylation modifying enzymes including DNMT3A and TET1, indicating their possible roles of recruiting these enzymes. Third, these motifs significantly overlap with SNPs associated with gene expression and those with DNA methylation. Fourth, disruption of these motifs by SNPs is associated with significantly altered methylation level of the CpGs in the neighbor regions. Furthermore, these motifs together with somatic SNPs are predictive of cancer subtypes and patient survival. We revealed some of these motifs were also associated with histone modifications, suggesting possible interplay between the two types of epigenetic modifications. We also found some motifs form feed forward loops to contribute to DNA methylation dynamics.

## Introduction

DNA methylation plays crucial roles in many biological processes and aberrant DNA methylation patterns are often observed in diseases. There are three DNA methyltransferases (DNMTs) in human that are responsible for *de novo* or maintaining methylation of cytosine. Although these enzymes themselves do not show strong sequence preference *in vivo*, DNA methylation is highly locus specific such as hypo-methylation of active promoters and enhancers. An urging question is how such locus-specific DNA methylation pattern is established. One of the possible mechanisms is that DNA binding proteins or non-coding RNAs recognize specific DNA motifs and their binding recruits DNMTs to a particular locus to methylate cytosines in the region. These factors can be specifically active in a cell type or state such that to provide the cell type-and locus-specificity. Accumulating evidences suggest that protein binding such as CTCF and other proteins can create low methylated regions in the regulatory sites and introducing specific nucleotide sequences can establish DNA methylation^1,2^. These observations suggested the importance of the DNA sequence in shaping methylation state. Several studies have illustrated the relationship between sequence features and DNA methylation^3–13^ but the DNA motifs recognized by the DNA methylation associated proteins have not been well characterized. Therefore, cataloging these motifs would pave the way towards understanding the mechanism of the locus-specificity of DNA methylation.

Cataloguing DNA methylation associated motifs requires a comprehensive set of methylomes and whole-genome bisulfite sequencing (WGBS) is a common technology to map DNA methylation in the entire human genome. The NIH Roadmap Epigenomics Project^14^ has generated WGBS data in 34 cell lines or tissues, which provides an opportunity to discern motifs associated with DNA methylation. We reason that contrasting regions that are commonly methylated across cells/tissues to those commonly unmethylated would increase the signal-to-noise ratio to identify the motifs most relevant to DNA methylation. Furthermore, to consider the impact of cell type and cell state on DNA methylation, we also need to uncover motifs associated with variable methylation levels across cells/tissues; a caveat is that these motifs can be confounded by those only related to cell specificity. To this end, we have defined commonly methylated (unmethylated) regions across the 34 cells (CMR/CUR) as well as variably methylated (unmethylated) regions (VMR/VUR) that show cell-specific methylation (unmethylation). We have found the DNA motifs that are discriminative of these regions.

To confirm the motifs’ association with methylation, we overlapped them with DNMT and TET ChIP-seq peaks and observed strong enrichment. We also used TCGA dataset to further assess the importance of these motifs in shaping DNA methylation. Interestingly, we found that, if these are SNPs occurring in the motifs, the methylation levels in the nearby CpGs are significantly altered, i.e. perturbation to a MM (UM) motif in a highly (lowly) methylated region would decrease (increase) the local methylation level. This observation strongly supports the functionality of the identified motifs in establishing or maintaining locus-specific DNA methylation. Furthermore, we observed eQTL and mQTL SNPs are enriched in the found motifs. We also found that the combination of somatic SNPs and the found motifs can significantly improve the prediction accuracy of cancer type and patient survival than using SNPs alone. This observation also supported the functionality of the DNA methylation associated motifs. Additional analyses also revealed the potential interplay between DNA methylation and histone modification as well as their contribution to DNA methylation dynamics.

## Results

### Defining DNA methylation regions and *de novo* motif discovery

We aimed to identify DNA motifs associated with DNA methylation and thus started with searching for methylation regions that have the strongest signals. We collected whole genome bisulfite sequencing (WGBS) data of 34 human methylomes generated by the NIH Roadmap Epigenomics Project^15,16^ (**Figure 1A**). We took an approach similar to the Ziller *et al.* study^17^ and defined 1.55 million methylation regions containing 11.5 million CpG sites in the 34 methylomes. Because the methylome data is noisy, we only considered regions containing 2 or more CpGs within 400 bp apart, which covers 29.2% of the human genome.

**Figure 1.**
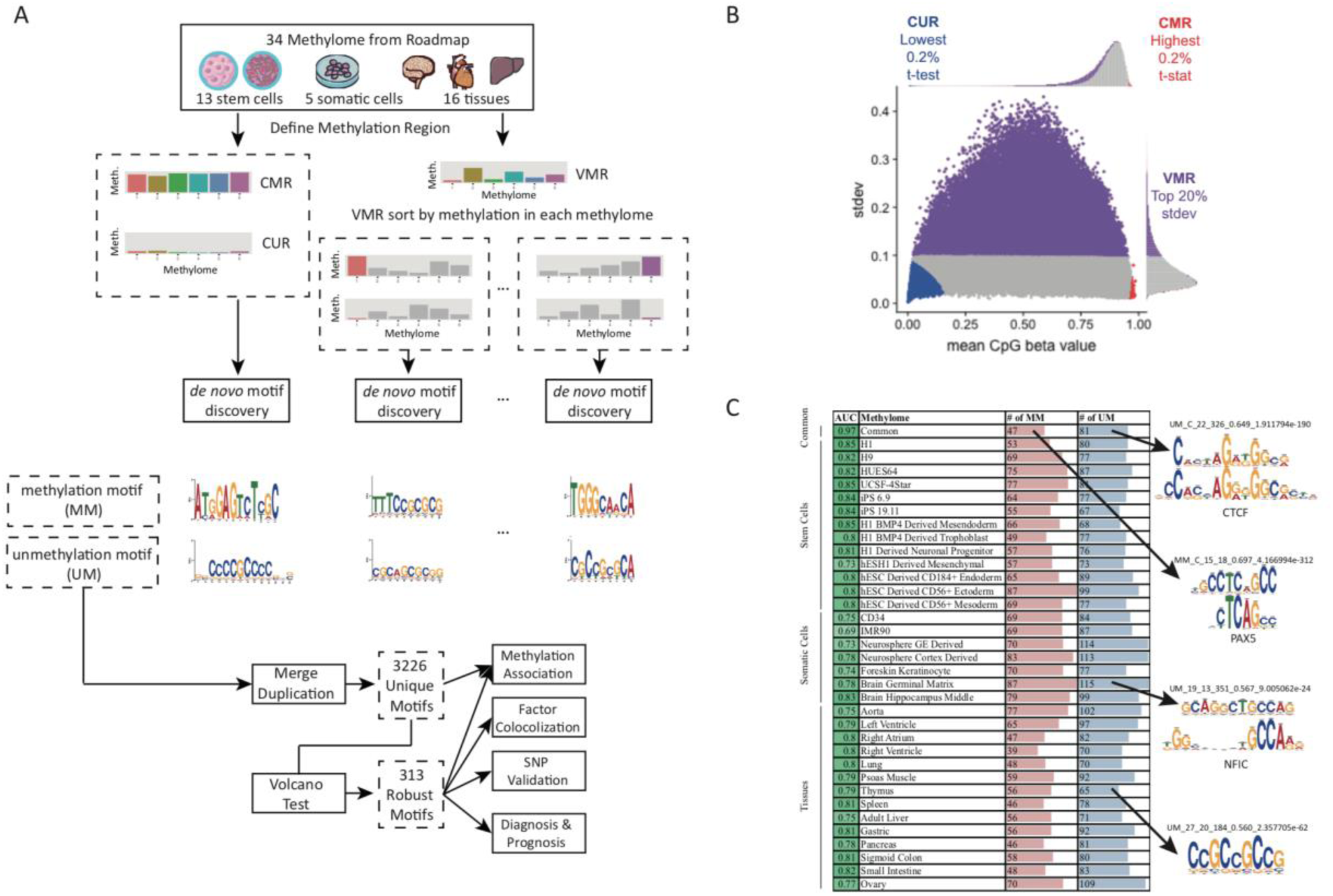
Defining methylated regions and searching for methylation associated motifs **A**. The strategy of identifying DNA methylation associated motifs. **B**. WGBS CpG sites are merged within 400bp regions. Based on average CpG beta values of the region, we defined commonly methylated (CMR), un-methylated (CUR) and variably methylated regions (VMR). **C**. Identification of DNA methylation associated motifs in 34 cells and tissues. Example motifs are shown on the right (if matched to a known motif, the known motif logo is shown on the top).

Methylation level is associated with different functions. For example, low methylated regions (LMRs) are important in hematopoiesis and leukemia development^18^, DNA methylation valleys (DMVs) are long hypomethylated regions involved in development and tissue-specific regulation^19,20^; focal hypermethylation and long range hypomethylation are found in cancer^21^; variably methylated regions (VMR) are associated with histone modification and enhancer^22^. In this study, we defined three types of methylation regions based on the mean and standard deviation of the CpG methylation level in each region (**Figure 1A, 1B**): (1) Top 0.5% (or 7726) commonly methylated regions (CMR) which have highest methylation level across 34 methylomes; (2) Top 0.5% (or 7726) commonly unmethylated regions (CUR) with lowest methylation levels; (3) Top 20% (or 309040) variably methylated regions (VMR) with highest standard deviation and this percentage is consistent with the previously reported 21.8% to 22.6% VMRs in the methylome^17,22^. We are aware that these regions can vary upon the data sets used to define them. Because the 34 methylomes are derived from diverse cells and tissues, we argue the derived motifs are still a reasonable starting point of revealing DNA binding proteins recruiting DNA methylation enzymes.

Defining commonly and variably methylated/unmethylated regions allow identification of motifs that are associated with DNA methylation independent of cell type or cell-type specific. CMRs and CURs are regions that show consistent methylation pattern across a diversity of 34 cells and tissues, and therefore they likely harbor motifs associated with methylation/demethylation in a cell type independent manner. GREAT^23^ analysis showed CMRs are strongly (*p* < 1e-30) linked to DNA repair and mitosis and are mostly (68%) found in introns (**Figure S1A**), where hypermethylation has been reported to confer enhancer-like activity in EGR2^24^ and mediate alternative splicing^25^. CURs prefer promoters (66%) associated with (*p* < 1e-30) cell differentiation, development and morphogenesis, indicating the important roles of demethylation in these processes^26,27^(**Figure S1A**). By contrasting CMRs to CURs, we identified 55 CMR and 87 CUR motifs using a motif finding algorithm Epigram^3^ (**Figure 1A, 1C**). A 5-fold cross-validation using Epigram^3^ successfully discriminated CMRs from CURs using the motifs (AUC= 0.97) (**Figure 1C**). Note that Epigram balances the GC content, sequence number and length in the foreground and background, which avoids identification of trivial sequence motifs (see details in **Methods** and ref. ^3^). Because these motifs are associated with high or low methylation regions commonly shared by a diverse cell types, it is reasonable to argue that they are important or even casual for establishing, maintaining or removing DNA methylation.

Similar to TFs whose binding motifs are defined but their activities are cell type specific, the usage of DNA methylation associated motifs is determined by cellular state. The VMRs show cell type specific methylation patterns, which provides an opportunity to identify motifs active in particular cell types. We contrasted top 6000 methylated and unmethylated VMRs sorted in each cell type and discovered average 63 methylation and 85 unmethylation associated motifs in each methylome, with an average AUC of 0.79 (**Figure 1C**).

In total, 5172 motifs were identified from 35 Epigram runs (1 common + 34 cell-specific). Because the same or similar motifs could be found in multiple cells, we clustered these motifs into 3226 unique ones using motif similarity measurement based on Jensen-Shannon divergence (see **Methods**). To control false discovery rate (FDR), we further conducted a robust volcano test^28^ with stringent *p*-value < 1e-10 and enrichment > 2 requirement, resulting in 313 methylation motifs for the follow-up analysis (**Figure 1A, S1B**), including 221 unmethylation motifs (UM) and 92 methylation motifs (MM). Among them, 36 (16.2%) and 14 (17.1%) are matched to 50 known motifs in the latest version of HOCOMOCO^29^. The matched included previously confirmed factors to influence methylation level such as CTCF^2^ and PAX5^30^ as well as factors KLF4, SP4, and EGR1 that have been reported to regulate gene expression by binding to CpG rich promoters^31^ (**Figure 1E**). The majority of the motifs are novel and showed strong sequence preference. UMs are more similar to each other and have higher GC content (eg. CCGCCGCCG) than MMs (**Figure S1C, S1D**). Note that these motifs were found by Epigram after sequence balancing which removes GC content bias^3^. While high GC content and CpG-rich sequences have been associated with hypomethylation in regions such as CG-islands^32^ and in specific cells^33–35^, our analysis revealed specific DNA motifs with sophisticate patterns that may be recognized by proteins or ncRNAs.

### Identified motifs are associated with the local DNA methylation deviated from the background

We first investigated the DNA methylation levels around the identified motif occurring sites (determined by FIMO^36^ using *p* < 10^−5^, the same parameters were used for all the relevant analyses thereinafter). We did observe hypomethylation and hypermethylation in the neighbor CpGs of the UM and MM motifs, respectively. Several representative examples are shown in **Figure 2A.** It is obvious that DNA methylation levels around the motif sites show a sharp “dip” or “peak”, suggesting the association is highly locus-specific. Interestingly, this trend remains the same in different cell types despite that the methylation levels in the surrounding regions vary. For example, motif UM_238.2_3.88_0.53_5 (matched to the WT1 motif) was identified from VMRs in the right ventricle tissues; the methylation level at its occurring sites decreases in all the cell types although the methylation level ranges from 0.6 to 0.8 in the surrounding regions (**Figure 2A**). This observation confirms the functionality of individual UM and MM motifs even though the local environment is overall hyper-or hypo-methylated.

**Figure 2.**
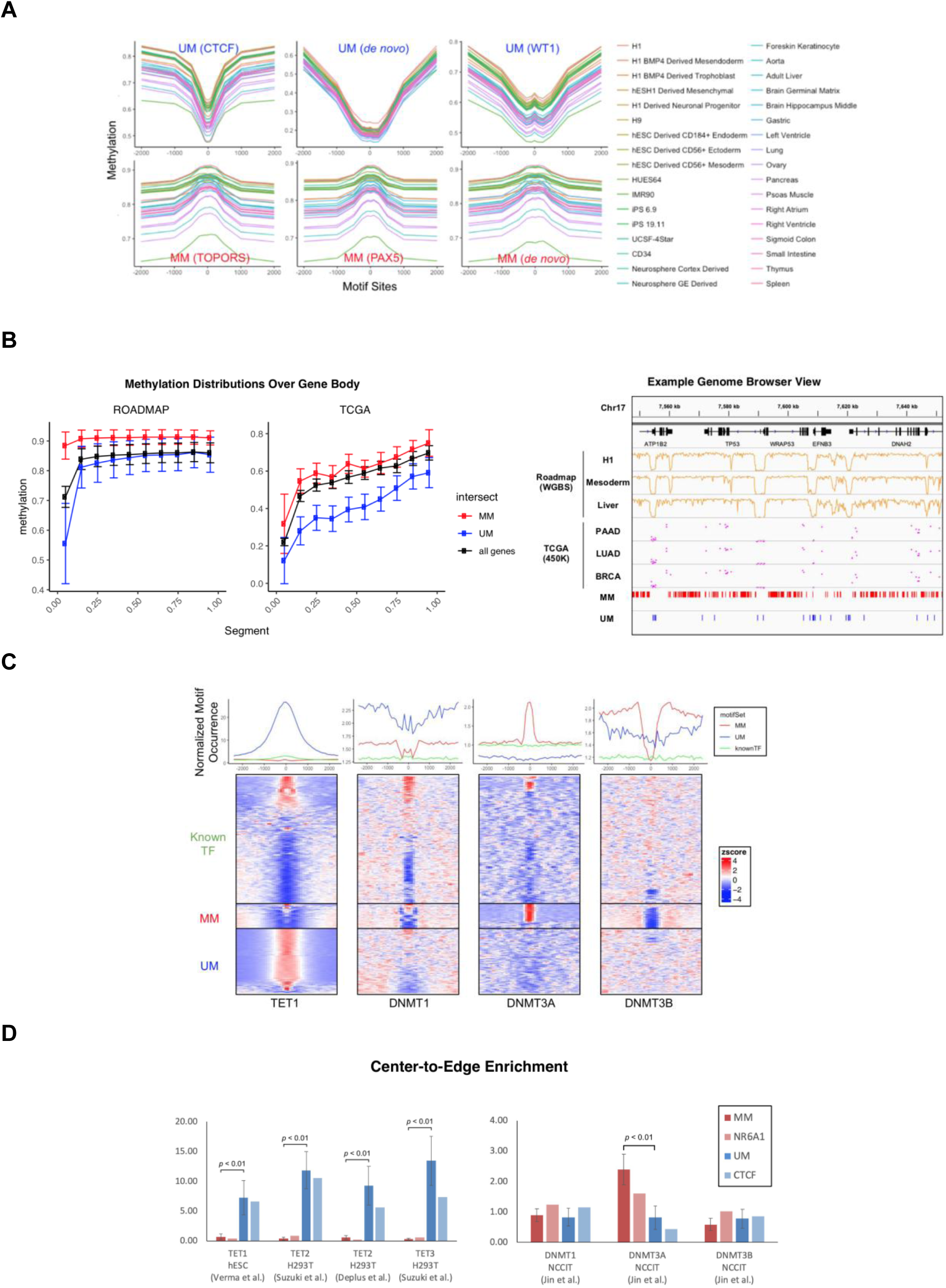
Identified motifs marks methylation level **A.** Example motifs shown with average CpG methylation level calculated in 50 bp bins around all motif sites, determined by FIMO at 1e-5 *p*-value cutoff. Upper panel, from left to right: UM_180.0_3.14 (matched to CTCF); UM_106.1_4.08 (*de novo*); UM_238.2_3.88 (matched to WT1); lower panel, from left to right: MM_65.9_2.90 (matched to TOPORS); MM_814.4_2.02 (matched to PAX5); MM_206.3_2.16 (*de novo*). **B**. DNA methylation levels in the ROADMAP (left) and TCGA (right) data sets over gene body. Each gene body was split into ten equal bins and the Beta values of all CpGs in the same bin were averaged over all genes. Lower panel shows the correlation between the motif occurrences and CpG methylation in ROADMAP (WGBS data from H1, mesoderm and liver) and TCGA (450K methylation of CpGs averaged in patients from PAAD, LUAD and BRCA) around TP53 (chr17:7,540,000 - 7,650,000) **C.** Normalized motif occurrence of UM, MM and known TFs (excluding matched) from HOCOMOCO^29^ at 5000 bp windows centering ChIP-seq peaks of TET1, DNMT3A and DNMT3B collected from various studies^45–47^. Lower panel shows the clustered heatmap of normalized z-score. **D.** Center-to-edge enrichment of UMs and MMs in comparison with TF NR6A1 and CTCF, which were reported to recruit DNMT and TET to specific loci, at the ChIP-seq peaks of DNMTs and TETs.

We further examined the impact of these motifs on methylation in the gene coding regions. UM and MM consistently mark lower and higher local CpG methylation levels in the gene coding regions (**Figure 2B**). In the Roadmap dataset, we observed a significant impact of UMs or MMs on DNA methylation level around the transcription start sites (TSS) (**Figure 2B, left panel**). DNA methylation in the promoters is important for regulating gene expression^37^ and thus itself is likely under active regulation. We observed the same trend in the TCGA DNA methylation data of 9037 patients from 32 cancers measured by Illumina 450K array^38^ (**Figure 2B, right panel**). On average, CpG methylation decreases from the beta value of 0.81 in the Roadmap dataset, dominated by normal cell lines and tissues, to 0.59 in the TCGA cancer patients across 20,260 protein-coding genes. This observation is consistent with the global hypomethylation in cancer cells that have been reported in the literature^19,26,39^. However, the MM and UM occurring bins still showed respectively higher and lower methylation levels than the background. As an example, UM and MM occurrence sites are characterized by lower and higher methylation in the gene coding region of TP53 (chr17:7,540,000 - 7,650,000) in both TCGA and Roadmap data. Collectively, our results on two separate data sets generated by different technologies support that the identified DNA motifs play critical roles in influencing the local CpG methylation.

### Identified motifs are significantly enriched at TETs and DNMTs binding sites

Locus-specific DNA demethylation or methylation depends on the recruitment of specific enzymes such as TET^40^ and DNMTs^41^ to particular genomic regions^42–44^. We reasoned that, if the identified motifs are important for recruiting the enzymes, these motifs would be enriched around the binding sites of the recruited enzymes. To this end, we have collected all the available ChIP-seq experiments of TET and DNMT enzymes^45–48^. Indeed, at the center of TET1 ChIP-seq peaks in hESC H1 cells^46^, the UM sites occur 26.7 times of expected counts (see details in **Methods**), whereas MM motifs occur roughly same (1.4 times) as the expected counts (**Figure 2C**, first panel from the left). This observation is consistent with the previous reports that TETs can be recruited to specific locus by DNA binding factors^40,44^. Interestingly, the wide distribution of UM around TET peaks compared to MM-DNMT overlap is consistent with the previously reported role of TET in protecting spanned low-methylation regions termed methylation canyons against hypermethylation^49^. Furthermore, TET prefers CpG-rich patterns such as CpG island which spans several kilobases^50^ and can bind CpG-rich DNA sequences^42^ in mammalians to maintain stable demethylation^51^; consistently, UMs have significantly higher GC content than MMs and known motifs (*p* < 0.05, **Figure S1C**).

We observed different motif occurring patterns around the binding sites of different DNMT enzymes. DNMT3A and DNMT3B are responsible for *de novo* methylation^52^. At the center of DNMT3A ChIP-seq peaks in the human NCCIT cells^45^, we observed a peak of the MM motif occurrence compared to the known and UM motifs (**Figure 2C**). Interestingly, the MMs are enriched at the shoulder regions of the DNMT3B binding sites but depleted at the center (**Figure 2C**). Note that only 2.2% of DNMT3A and 3.8% of DNMT3B peaks overlap with each other^45^ (**Figure S2A**). Several studies have demonstrated some distinct roles of DNMT3A and DNMT3B, showing that DNMT3B preferentially targets gene bodies marked with H3K36me3^53–56^; in fact, H3K36me3 is 4.27 times enriched at the DNMT3B compared to DNMT3A peaks in gene coding regions (**Figure S2A**). These observations suggest that the MMs are likely recognized by DNA binding factors involved in actively recruiting DNMT3A, whereas DNMT3B may be recruited by flanking sequences containing MMs and together with chromatin marks and/or other factors such as H3K36me3. Interestingly, DNMT1, an enzyme involved in DNA methylation maintenance and recognizing hemimethylation^57^, shows a different profile from DNMT3A/B (**Figure 2C**, second panel from the left). This difference may be resulted from the different mechanisms or factors involved in active and passive DNA methylation.

To further validate if the observed co-occurrence around methylation enzyme is significant, we also compared the center-to-edge enrichment of UM and MM with TFs known to regulate DNA methylation (**Figure 2D, method**). Previous studies have reported that introducing a CTCF binding site at a particular locus leads to TET recruitment and local DNA demethylation^2^. NR6A1 has also been confirmed to recruit DNMT to methylate at target genes^58^. Here, we show that at the center of TETs binding sites, UMs are significantly more enriched than MMs, and have even higher enrichment than CTCF (**Figure 2D**, left panel). Similarly, MMs are significantly more enriched than UMs at the center of DNMT3A binding sites, surpassing that of NR6A1 (**Figure 2D**, right panel). The enrichment of MMs and UMs were further compared with the known TFs such as PAX5, TOPORS, WT1 and PPARG that are most enriched at the TETs and DNMT3A sites (**Figure S2B**). These results demonstrated that the identified motifs can be recognized by particular DNA binding factors that in turn recruit the methylation modifying enzymes in a locus-specific manner. Given that the majority of MMs (71.4%) and UMs (83.9%) are *de novo* motifs, our findings pave the way towards identifying particular factors involved in locus-specific methylation regulation.

### SNPs occurring at identified DNA motif sites is associated with altered methylation level

To validate the functionality of the identified motifs, we investigated enrichment of functional SNPs (eQTL and mQTL) at motif occurrence sites. We analyzed the relationship between somatic mutation and methylation level using the TCGA data^38^ and identified methylation quantitative trait loci (mQTL), which are SNPs correlating with CpG variation within 5000bp. Using Matrix eQTL^59^, we identified 26341 SNP-CpG pairs (mQTL), corresponding to 17038 unique SNPs and 20043 CpGs, from the total 1.3 million somatic mutations in 9037 patients of 32 cancers. We observed an average 11.7% mQTL discovery rate at the motif sites compared to 2.3% in the background (**Figure 3A, upper left panel**). This enrichment difference is most prominent around transcription start site, suggesting that the identified motifs have stronger impact on methylation at TSS (**Figure 2B**)^60–62^. Enrichment of mQTL in both MM and UM sites was also found in three additional human methylome datasets using the reported mQTLs in the original studies^63–65^ (**Figure S3A**), which confirms the generality of this observation. Because DNA methylation is associated with gene expression^17,66^, it is not surprising that MMs and UMs significantly overlap with expression quantitative trait loci (eQTL), which are SNPs correlated with gene expression level (**Figure 3A, right panel**).

**Figure 3.**
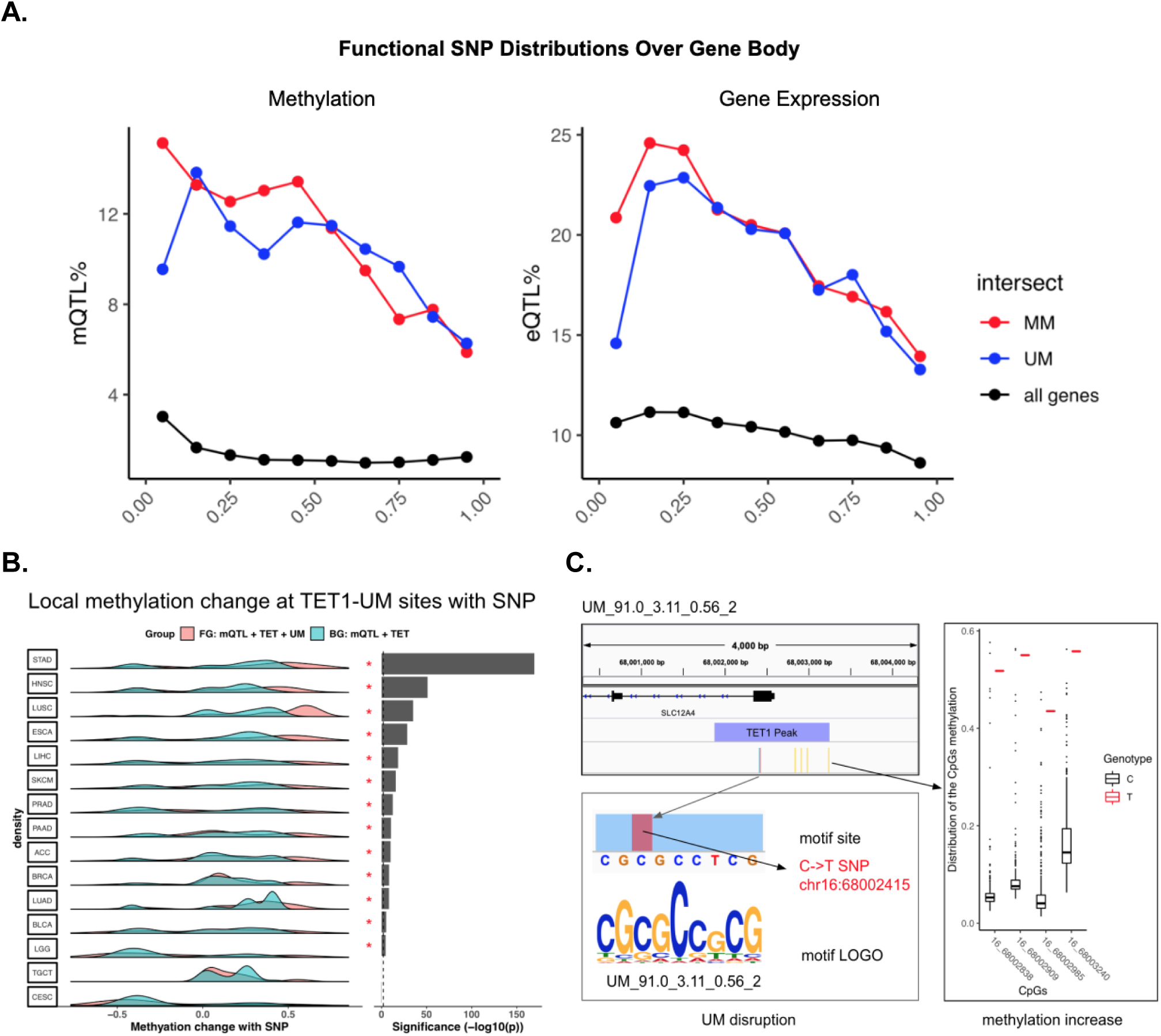
SNP at motif sites co-occur with local methylation alteration **A**. Distribution of mQTLs (SNPs associated with methylation) and eQTLs (SNPs associated with gene expression) over gene body (see **Methods** for details). Each gene body is split into ten equal bins. **B.** Methylation level change of CpG sites nearby TET1-UM sites (TET1 binding peaks containing UM motifs) overlapping with somatic SNPs. Asterisks indicate *p* < 0.01 calculated with paired one-tail t-test, pairing foreground observed methylation change to the corresponding background expected methylation change. Foreground (FG), mQTL at TET1-UM sites. Background (BG), mQTL at TET1 binding peaks^45–47^. To ensure the statistical significance, we only considered the cancers with >100 CpGs within 5000bp of TET1-UM sites (see details in **Methods**). **C.** An example showing disruption of an UM motif (no match with known motifs) by a C->T somatic mutation at chr16:68002415 significantly increases the methylation level of the nearby CpGs in the LUAD patients.

To investigate the causality between these motifs and DNA methylation level, we analyzed whether disrupting these motifs would lead to DNA methylation change. We chose to focus on the possible binding sites of TET1 and DNMT3A containing these motifs because the significant enrichment of the found motifs in the enzyme-binding regions implies that the active methlyation or demethlyation is most likely mediated by DNA binding factors to recruit DNMT3A or TET1, respectively. Despite the ChIP-seq experiment of TET1/DNMT3A was done in one particular cell type, the sequence features, i.e. the motif composition in these regions, do not change and thus the mechanism of the active methylation regulation. The methylation change is decided by which factors are expressed and active in a specific cell type or state. Disrupting these motifs would lead to methylation change in the nearby CpGs.

Using the TCGA data, we first identified 5372 CpG sites from 15 cancers within 5000bp of the TET1 binding peaks that also contain SNPs overlapping with UMs in at least one patient. Because we did not have TET1 ChIP-seq data in the cancer patients, we used the published data measured in hESC (see **Figure 2C/D**). We compared the methylation change of these CpGs between patients with and without SNP in each cancer. 13 out of 15 cancers showed significant (*p* < 0.01) increased methylation level of with-SNP compared to the without-SNP patients (background) (**Figure 3B,** see **Methods** for details). One example is given in **Figure 3C** for an UM motif UM_91.0_3.11_0.56_2. This motif is within a TET1 peak and is disrupted by a C->T somatic mutation at chr16:68002415 on the first exon of SLC12A4 in one LUAD cancer patient. All 4 CpGs within 500 bps upstream of the SNP showed increased methylation (beta value increased from 6.2% to 52%, 8.8% to 55%, 6.2% to 44% and 17% to 56%, respectively). Hypomethylation in the SLC12A4 promoter is related to resistance to platinum-based chemotherapy in ovarian cancer^67^; the 4 CpGs affected by the SNP are located in the SLC12A4 promoter, suggesting a mechanism of how the SNP may affect response to chemotherapy through regulation of local DNA methylation. More examples of SNP-induced methylation change through disrupting UMs are shown in **Figure S3B.**

Overlapping MM and SNPs with DNMT3A peaks only resulted in <100 CpGs sites in 2 cancers. Although we observed decreased methylation level of DNMT3A-MM overlapping with mQTL as predicted, the analysis did not have enough statistical power. Because the methylation was measured by 450K array and SNPs were detected by Affymetrix Genome-Wide Human SNP Array 6.0, it is reasonable to expect that more sites can be observed with whole methylome and whole genome sequencing data.

### Combining Motifs and SNPs Shows Diagnosis and Prognosis Power

DNA methylation has been shown to be predictive for cancer diagnosis and patient survival prospective^68,69^. Since we have shown motif disruption is associated with methylation change, we hypothesized that combining motifs with SNPs can improve prediction for cancer diagnosis and patient survival. To evaluate this, we trained gradient boosting models^70^ using SNP and SNP+motif as features in 32 TCGA cancers from 7120 patients (see **Methods** for details). We calculated both auROC and auPRC (a metric for imbalanced dataset to avoid inflated evaluation of the performance)^71^. Inclusion of the motifs in the models showed increased auROC and auPRC in all the 32 cancers. On average, auROC increased from 0.78 to 0.92 and auPRC from 0.45 to 0.56, whereas 26 (for auROC) and 13 (for auPRC) improvement are statistically significant (*p* < 0.01) (**Figure 4A**). Notably, several cancers showed drastic improvement, including ovarian cancer (OV, auPRC from 0.41 to 0.79), thyroid carcinoma (THCA, auPRC from 0.49 to 0.82), acute myeloid leukemia (LAML, auPRC from 0.6 to 0.88), pheochromocytoma and paraganglioma (PCPG, auPRC 0.49 to to 0.75) (**Figure S4B**). These cancers all have reported aberrant methylome and have methylation associated diagnosis and therapeutic targets^72–75^.

**Figure 4.**
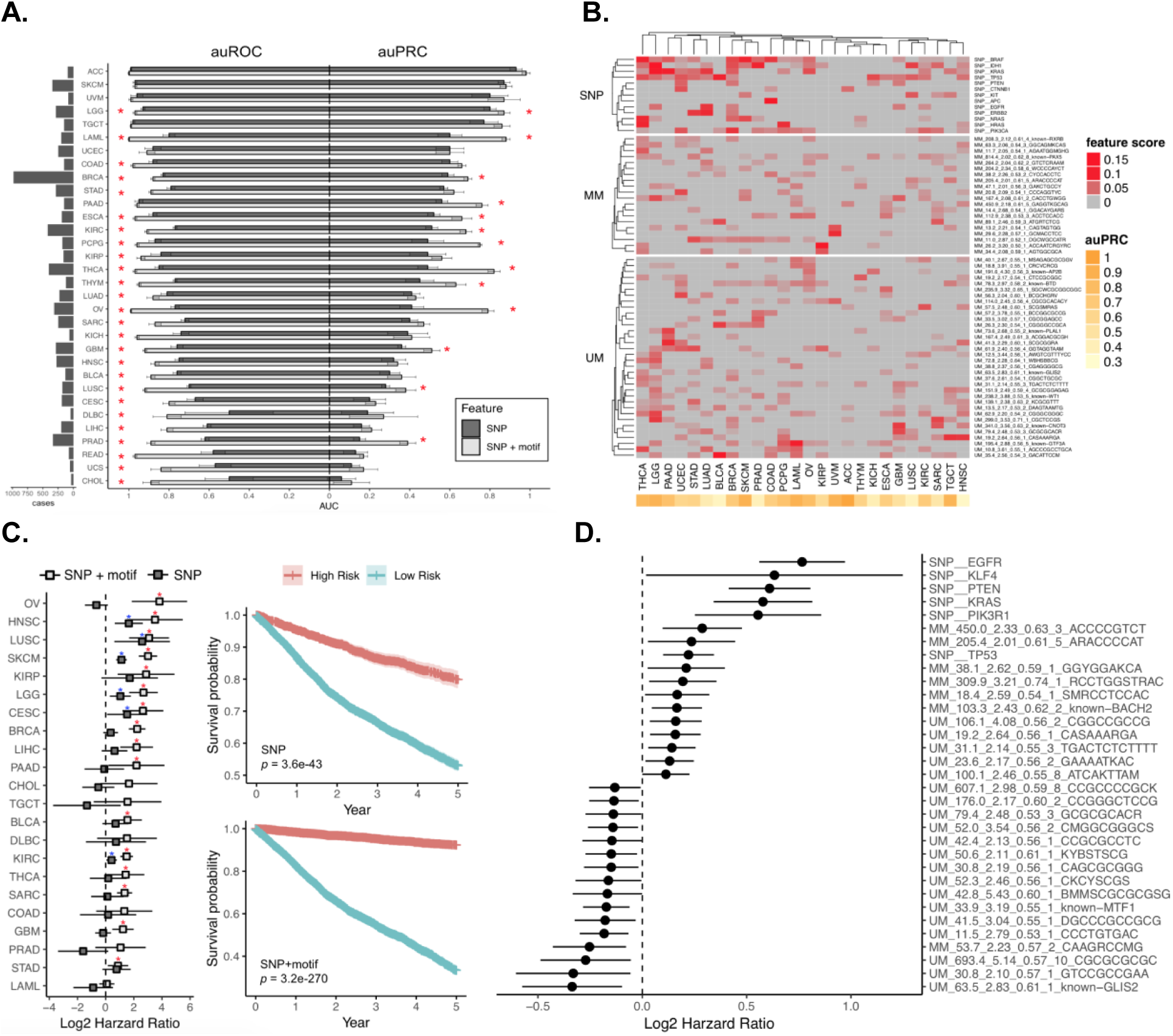
Motifs disrupted by SNPs predicts cancer subtype and survival **A**, auROC and auPRC for cancer subtype prediction. Classification model of each cancer built with gradient boosting. Performance evaluated with auROC (area under receiver operating characteristic, good for overall evaluation.) and auPRC (area under precision recall curve, good for unbalanced dataset where positive label is scarce). SNP: using somatic mutation as features. SNP+motif: using both somatic mutation and collective disruption of motif site as features (See **Methods** for details). * Adjusted *p* < 0.05. **B**, Results of top predictive features (score>0.01) using gradient boosting out-of-bag estimation. 26 cancers with auPRC > 0.3 are shown. **C**, Results of survival analysis with gradient boosting model using SNP or SNP+motif as features. Left: multivariate survival analyses for all solid TCGA cancers. Forest plots showing log2 hazard ratio (95% confidence interval) of predicted high risk group by SNP+motif or SNP. *Adjusted *p* < 0.05 (blue for SNP and red for SNP+motif). Right: Kaplan-Meier survival estimation (95% confidence interval) in high-risk group versus low-risk group predicted by SNP (top) or SNP+motif model (bottom). **D**. Multivariate survival analysis showing factors correlating with patient survival (*p* < 0.05) with log2 hazard ratio (95% confidence interval).

For 26 cancers with auPRC>0.3, the 67 most predictive features (score > 0.01) determined by the gradient boost estimator are shown in **Figure 4B (**see **Methods** for details), including 13 SNPs, 20 MMs and 34 UMs. Only 2 MMs are matched to known motifs (RXRB and PAX5), whereas 7 UMs to AP2B, BTD, PLAL1, GLIS2, WT1, CNOT3 and GTF3A. The predictive SNPs include those occurring on the cancer driver genes such as BRAF (in 16 cancers), TP53 (in 14 cancers), IDH1 (in 14 cancers), PIK3CA (in 13 cancers) and KRAS (in 12 cancers). Strikingly, we found numerous MMs and UMs very predictive in multiple cancers. Notably, MM_814.4_2.02_0.62_8 (PAX5) that has been shown to strongly impact local methylation level (**Figure 3C**) is important in 12 cancers. The 5 UMs predictive in >10 cancers are UM_78.3_2.97_0.58_2 (BTD), UM_13.5_2.17_0.53_2, UM_195.4_2.88_0.56_5 (GTF3A), UM_35.4_2.56_0.54_3 and UM_61.9_2.40_0.56_4 (**Figure 4B**).

To evaluate the prognosis power of the motifs, we trained two gradient boosting models (SNP and SNP+motif) to discriminate low-risk from high-risk patients. We evaluated the performance using survival hazard ratio of the predicted high-risk group (higher ratio means better performance). The SNP-only model found 6 out of 22 cancers having significant (*p* < 0.05) hazard ratio. In comparison, SNP+motif model achieved 16 out of 22 cancers having significant (*p* < 0.05) hazard ratio (**Figure 4C, left panel**, see **Methods** for details). Kaplan-Meier test showed better separation of patient survival between the predicted low-risk and high-risk groups by considering motifs (*p*=3.6E-43 for SNP and *p*=3.2e-270 for SNP+motif, **Figure 4D, right panel**). Multivariate survival analysis on the full model revealed important factors correlated with patient survival (*p* < 0.05), including 6 SNPs, 6 MMs and 20 UMs (**Figure 4D**). These results further confirmed the functionality of the discovered motifs and highlighted the potential for clinical application.

### Motifs involved in both DNA methylation and histone modifications

Both DNA methylation and histone modification play important roles in regulating gene expression and their interplay has been well recognized^76,77^. In a separate study, we identified 361 motifs that are associated with 6 (H3K4me1, H3K4me3, H3K27ac, H3K27me3, K3H9me3, H3K36me3) histone modifications from 110 diverse human cell types/tissues^78^ (**Figure 5A**). By comparing the 313 methylation motifs and the histone associated motifs, we found that 56.5% MMs (52 out of 92) overlap with histone motifs (e-value cutoff of 0.05 using TomTom). Unsurprisingly, 35 MMs are aligned to H3K36me3 motifs as H3K36me3 can recruit DNMT3A/3B through their PWWP domain^79,80^. In contrast, 74.2% (164 out of 221) UMs found no match to histone motifs. 57 UMs are matched to motifs associated with active promoter or enhancer marks: 12 UMs matched to H3K27ac, an active promoter and enhancer mark; another 12 UMs matched to the promoter mark H3K4me3. As active enhancers and promoters tend to have low methylation^15^, this observation is not unexpected. Interestingly, we observed another 12 UMs matched to the motifs associated with the poised promoter markers H3K4me3+H3K27me3. Previous studies also suggested colocalization of H3K4me3 and H3K27me3 marks is associated with DNA hypomethylation in pre-implantation embryos^81^.

**Figure 5.**
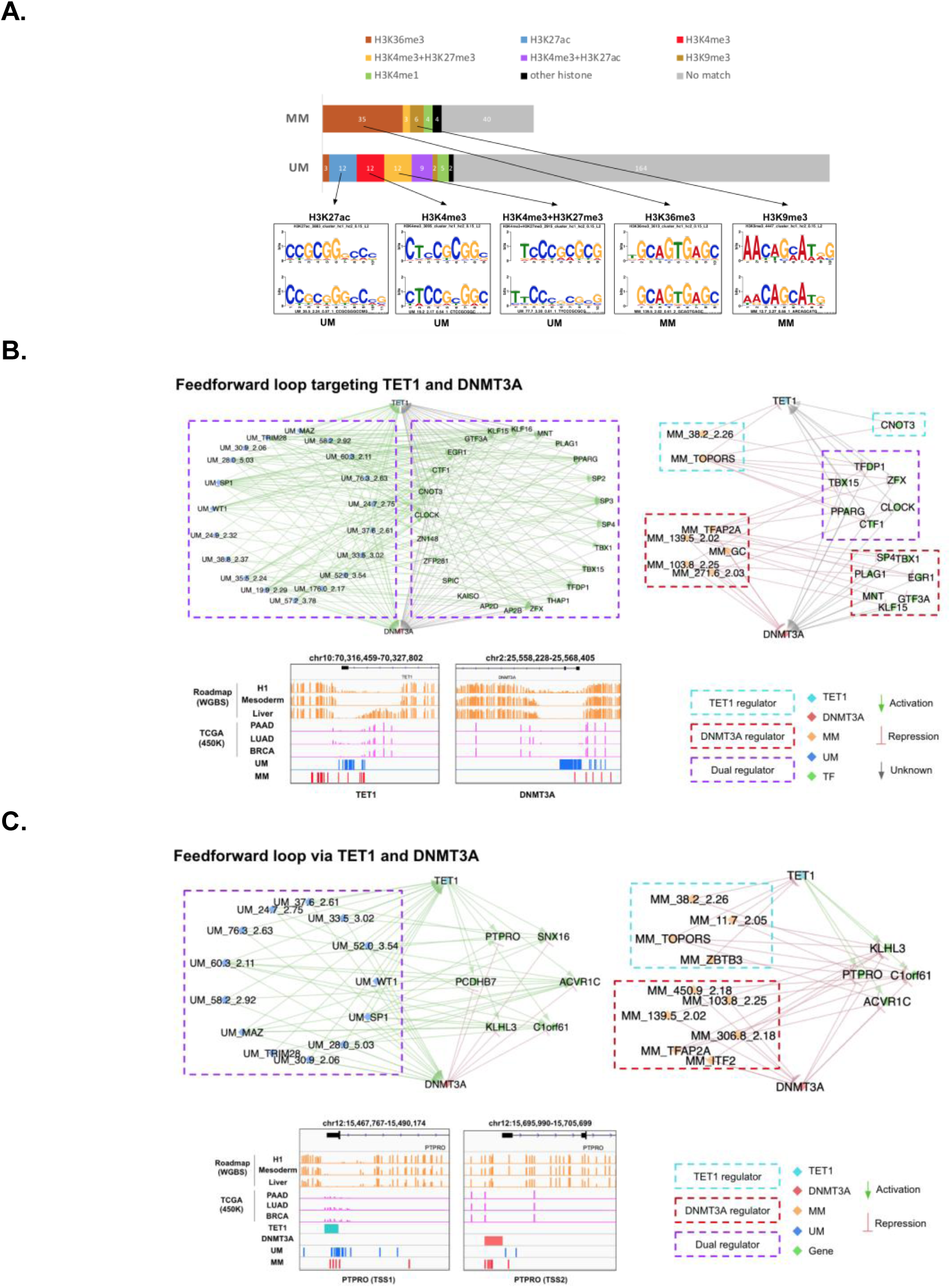
Methylation motifs interplay with TET1, DNMT3A, gene regulation and histone modification **A.** Methylation motifs matched to histone motifs^78^. Motifs are aligned with TomTom with e < 0.05. Lower panel showing several examples. **B.** Feedforward loop targeting TET1 and DNMT3A. **C.** Feedforward loop via TET1 and DNMT3A.

### Regulatory loops on DNA methylation

DNA methylation is dynamically regulated in response to the cell state change. We analyzed the putative regulatory connectivity between the identified motifs, transcription factors and the modifying enzymes of TET1 and DNMT3A. We only considered TET1 and DNMT3A here because their binding peaks are significantly enriched with UMs and MMs, respectively (**Figure 2C**). It is well accepted that a known TF motif occurring in the promoter of a gene suggests a possible regulation of the gene expression by the TF. Similarly, we infer the occurrence of an UM or MM in a gene’s promoter indicates putative regulation on the DNA methylation level and thus affecting gene expression.

We first analyzed the promoters of TET1 and DNMT3A. We found 19 UMs in the promoters of both TET1 and DNMT3A. We also found these UMs appearing in the promoters of 25 TFs that also have motifs in the promoters of both TET1 and DNMT3A and presumably regulate the two enzymes (**Figure 5B**). Such a topology forms a feed forward loop (FFL)^82^ that involves three nodes: two regulator nodes (motifs and TFs), one regulates the other (motifs regulates TFs), and both jointly regulating a target (TET1 or DNMT3A) (see **Methods**). UMs induce demethylation of TET1/DNMT3A and their regulator TFs, which forms positive FFLs to enhance the expression of both TET1 and DNMT3A once the motifs are activated. We also found 2 and 5 MMs occurring in the promoters of TET1 and DNMT3A, respectively. These MMs appear in the promoters of 14 TFs as the other regulator of TET1 or DNMT3A, of which 1 TF only regulates TET1, 7 TFs only regulates DNMT3A and 6 TFs regulate both (**Figure 5B**); these FFLs form enhanced dynamic regulation to repress TET1 and DNMT3A expressions. Overall, there are many more activating than repressive FFLs on regulating TET1 and DNMT3A.

Previous reports have also shown TET1 and DNMT3A have competitive binding to regulate promoters in mouse embryonic stem cells^83^. In addition, in honey bee Dnmts and Tet (homolog of vertebrate DNMTs and TETs) were found to target memory-associated genes sequentially, while Dnmt3 was found in a negative feedback loop for DNA methylation^84^. We found 6 genes targeted by UMs and also by both TET1 and DNMT3A (as indicated by their ChIP-seq peaks in hESC and NCCIT cells, respectively) (**Figure 5C**). Interestingly, 4 of them (KLHL3, C1orf61, ACVR1C, PTPRO) are also targeted by MMs and either TET1 or DNMT3A (**Figure 5C**). One of them, PTPRO, a cancer suppressor and therapeutic target of a variety of solid and liquid tumors, is silenced by promoter hypermethylation^85^. In fact, we observed higher methylation at the promoter of the first TSS of PTPRO (TSS1, chr12:15,474,979-15,476,332) in the TCGA patients (beta value average at 0.15) compared to the ROADMAP methylomes (beta value averaged at 0.05) (**Figure 5C**). PTPRO has multiple TSSs and alternative splicing forms^86^, and each TSS has a TET1 or DNMT3A ChIP-seq peak (**Figure 5C**). As competitive binding of activator and repressor can lead to sharp turn on/off of the gene expression^87–89^, we speculate the competitive FFLs formed by the motifs and modifying enzymes would thus allow dynamic regulation of the methylation and presumably the expression levels of these genes.

## Conclusion

In this study, we present a comprehensive catalog of the DNA motifs associated with DNA methylation. We did observe coincident higher and lower methylation levels around the MM and UM occurring sites, respectively. Furthermore, the motif sites are also enriched with functional SNPs, including mQTL and eQTL. We also showed that combining DNA motifs and SNPs can achieve accurate prediction in TCGA cancer patient’s diagnosis and prognosis, which supports the importance of these motifs.

Our analysis suggested that these motifs are most likely involved in recruiting TET and DNMT3A/3B for active demethylation and methylation, as indicated by their significant enrichment in the binding sites of these enzymes. The passive or maintenance methylation mediated by DNMT1 seems to be regulated by mechanisms other than DNA binding co-factors because we did not observe enrichment of the found motifs in the DNMT1 binding sites. Interestingly, some of these motifs may also play roles in histone modifications as they were also found associated with histone modifications, particularly those relevant to DNA methylation such as H3K36me3 that were reported to recruit DNMT3A/B through their PWWP domains. Furthermore, these motifs can form feed forward loops (FFLs) with TFs to regulate TET1 and DNMT3A or regulate genes together with TET1/DNMT3A. These FFLs allow possible regulation of the DNA methylation dynamics and presumably the gene expression dynamics. Our motif analysis suggests putative mechanisms for experimental test.

## Methods

### *De novo* motif discovery

11.5 million CpG sites common across all human 34 methylomes have been collected from NIH Roadmap Epigenomics Project^15^. Methylation regions are defined by segments with 2 or more CpGs within 400 bp apart and region methylation level is defined by the mean CpG beta values. Each region is then assigned mean and standard deviation of methylation across all 34 tissues and cells. A t-test is done for all regions with the null hypothesis that a region is uniformly distributed from all CpGs in all tissues. CMRs and CURs are defined by highest and lowest 0.2% t-test scores, while VMRs are defined by the top 20% standard deviation (**Figure 1A, 1B**). For common motifs MM and UM, we perform Epigram contrasting CMRs and CURs.

In short, Epigram looks for enriched motifs that best differentiate the foreground from the background sequences. In both sets of the input sequences, Epigram iterates through all possible *k*-mers to calculate their occurrences, enrichment over genomic background and enrichment over shuffled input. These values are combined to determine the enrichment of *k*-mers. Position weight matrices (PWMs) are then generated by first picking a top *k*-mer and enriched *k*-mers similar to itself to construct a “seed” PWM, which is then extended by adding more enriched *k*-mers that are a few base pairs shifted from the original one. The motifs are then further ranked and filtered based on how well they differentiate the foreground from the background using LASSO (least absolute shrinkage and selection operator) logistic regression. The final set of motifs is then evaluated by random forest.

For tissue-specific VMM and VUM, we contrasted top 6000 most methylated and unmethylated regions in each methylome. In total, we identified 5172 motifs from 35 Epigram runs (34 methylome + 1 common) with default parameters^3^ before curation (**Figure 1C**). For each run, Epigram found DNA motifs that discriminate enrichment peaks of the high methylation region under consideration (eg. CMR) from a background of low methylation region (eg. CUR). Importantly, the background has the equal GC content, number of regions and sequence lengths as the foreground to avoid inflated prediction results caused by simple features or unbalanced data set.

### Motif curation and defining motif occurrence site

Following our previous study^3^, we match motifs to the 1156 known motifs documented by the HOCOMOCO ChIP-seq consortium^29^ using an e-value cutoff of 0.05 with Tomtom^90^. Next, we merged the similar motifs to remove redundancy. We calculated a pairwise motif distance using weighted Jensen-Shannon Divergence:

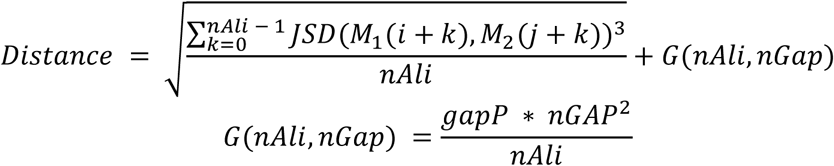

where *M*_1_, *M*_2_ are PWMs of the two motifs, respectively, *M(i)* represents the *i*th column in the matrix, *JSD(x,y)* is Jensen-Shannon divergence, *nAli* and *nGap* are respectively the length of the aligned sequence and gaps. Gap penalty function *G* has *gapP* as weight parameter set at 0.1. To ensure high similarity within motif cluster, gap penalty function is set to quadratic which is more stringent compared to traditional linear function to prevent having excessive gaps and hangovers. Motifs were hierarchically clustered with UPGMA^91^ algorithm and clusters were chosen using a distance cutoff of 0.1. As a result, we obtained 3226 clusters and selected the motif closest to the centroid of the cluster to represent all the motifs in that cluster. We combine the *p*-value of motifs in the cluster using Fisher’s combined probability test. Enrichment of each cluster is combined by geometric mean. The strongest 313 motifs are filtered by volcano test with combined *p* < 10^−10^ and enrichment > 2 (**Figure S1B**). Finally, motif occurrence sites are determined by a *p* < 10^−5^ calculated by FIMO^36^.

### Normalized motif occurrence and center-to-edge enrichment at DNMTs and TETs ChIP-seq peaks

DNMTs and TETs ChIP-seq peaks were downloaded from the published studies^45,47,48^. The 5000bp neighbor regions around the ChIP-seq peaks were included as the background or edge. Normalized motif occurrence was calculated using the following formula.

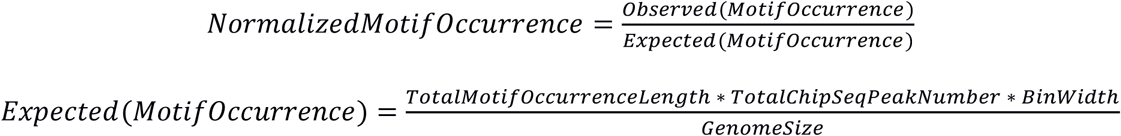

where *Motifsite count* is the occurrence number of a motif in a 100 bp bin, *Total Motif Occurrence Length*is the total length of genome-wide motif occurrences defined by FIMO (see the above section), *Total Chip Seq Peah Number*is the total number of ChIP-seq peaks, *BinWidth* is 100 bp and *Genome Size* is the genome size of 3.14E9 bp for the human genome hg19. We did this calculation for each of the 313 motifs in each 100 bp bin. Results are plotted in **Figure 2C**.

Further, center-to-edge enrichment was calculated by *Normalized Motif Occurrence* in the center 100 bp ChIP-seq bin divided by the average of *Normalized Motif Occurrence* at the bins 2500 bp upstream and downstream. Average enrichment and standard deviation were calculated across all MMs or UMs, followed by a two-tailed two-sample t-test, with *p* < 0.01 marked as significant. Results are plotted in **Figure 2D**.

### Functional SNPs enrichment analysis with TCGA

We downloaded the processed data (level 3) of 36 TCGA cancers from the Firehose database^92^ including patient survival, somatic mutations, 450K methylation array and RNA-seq data. Matrix eQTL^59^ linear model was used to identify mQTL and eQTL co-variating with methylation and transcript RNA-seq level, with 5000 bp distance cutoff from SNP to CpG and transcript TSS, respectively. We used a conservative *p*-value cutoff of 0.01 on top of a FDR cutoff of 10%. Then we calculated the number of mQTL or eQTL out of all SNPs in 10 bins of gene body, i.e., 0-10%, 10-20% … 90-100% of the mRNA transcript length, defined in Gencode v19^93^. We performed such analysis on all genes and repeated it with the UM and MM occurrence sites (**Figure 2A)**. To determine the significance of functional SNP enrichment, a chi-square test was carried out in each of the 10 bins of gene body, with the null hypothesis that mQTL% or eQTL% occurring at motif sites are the same as the rest of all genes, *p* < 0.01 are marked as significant.

### mQTL enrichment analysis with three independent datasets

Three human methylome studies with independently called mQTLs were collected, i.e. human life course study^63^, GenCord Cohort study^64^ and a Schizophrenia study^65^. In total, there are around 16,000 to 30,000 identified mQTLs collected from these published studies. We defined an enrichment score using the following formula.

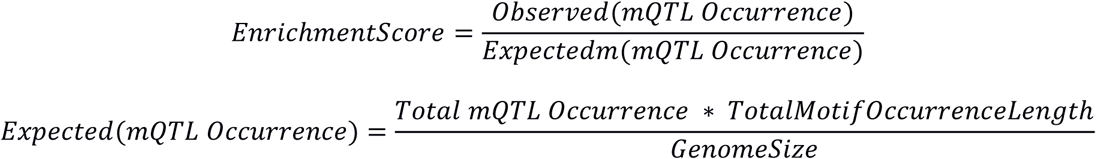

where *Total mQTL Occurrence* is the total number of mQTLs identified in each study, *Total Motif OccurrenceLength* is the total length of genome-wide motif occurrences defined by FIMO (see the above section) and *Genome Size* is the genome size of 3.14E9 bp for the human genome hg19.

We repeated this process in all samples from all three studies and calculated standard deviation. Specifically, (1) 5 life stages from birth, childhood, adolescence, pregnancy and middle age in human life course study (blood samples from 1018 mother–child pairs), (2) 3 tissues from fibroblasts, LCLs and T-cells in GenCord cohort by Maria *el al* (204 newborn umbilical cord samples) and (3) 3 regions from prefrontal cortex, striatum and cerebellum of adult brain regions in the Schizophrenia study (173 fetal brain samples ranging from 56 to 169 days post-conception). Finally, we used a single-tail one-sample t-test to determine the statistical significance (*p* < 0.01, **Figure S3A)**.

### Predicting TCGA cancer type with SNP and motif

For each of the 32 TCGA cancers (in total 7120 patients), we trained two gradient boosting models^70^ (SNP and SNP+motif) to distinguish a specific cancer from the other cancers. We chose gradient boosting implemented in sklearn^94^ and tuned its parameter based on a recent study^95^, which showed that this decision tree based model is robust and performs well. Note that TCGA has 4 aggregated cancer types (GBMLGG, COADREAD, KIPAN and STES) that combine individual cancers such as GBMLGG combining GBM and LGG; we excluded them from the 32 TCGA datasets to avoid inflating the performance due to using the same patients in both the training and testing sets. In a SNP-only model, the cancer subtype of each patient was predicted only by somatic mutations as features. Because the input features are large (1.3 million unique somatic mutations for 7120 patients), we first reduced feature number. Each feature was assigned a score by the gradient boosting out-of-bag importance and averaged in 5-fold cross validation to avoid overfitting. Features with negative importance scores were removed. Optimal number of features were determined as we observed the best model performance at around 500 features (**Figure S4A, upper panel**). Top 500 SNPs ranked by the average score were used, while assuring equal or better performance compared to the full model (**Figure S4A, lower panel**).

After feature selection, we obtained 500 selected SNPs. To represent what SNPs were found in each patient, we used a series (length 500) of 0s and 1s to indicate which SNPs a patient has. For example, 1,1,0,1, … indicates patient have the 1st, 2nd and 4th SNP. For a SNP+motif model, each patient was represented not only by these 500 selected SNPs, but also by whether each of the 313 motifs is disrupted by SNPs. We used a series (length 313) of integers to indicate how many SNPs (without feature selection) are harbored in the occurrence sites for each of the 313 motifs. For example, 10, 20, 0, … indicates there are 10 SNPs in all the occurrence sites of the 1st motif, 20 SNPs in the 2nd motif and no SNPs in the 3rd motif. The performances of the two models were evaluated by auROC and auPRC with 5-fold cross-validations for each cancer (**Figure 4A**). Feature importance was determined by the default out-of-bag (OOB) important scores using mean decrease of Friedman squared error over all cross-validated predictions in SNP+motif models. We filtered features with importance score >0.01 within the enriched 313 motif groups and well-studied SNPs located in the identified driver mutations by IntOGen Consortium^97^. To reduce false positives of selecting predictive features, we only considered 26 out of 32 TCGA cancers that showed auPRC > 0.3 (**Figure 4B**).

### Predicting TCGA patient survival with SNP and motif

All patients in 22 TCGA cancers with patient survival and SNP information were dichotomized based on 5-year survival to train two gradient boosting models (SNP and SNP+motif). We used the 500 SNP features and 813 SNP+motif features from the diagnosis analysis and cross validations were performed the same way as described above. The model performance was evaluated by the log2 hazard ratio and Kaplan-Meier estimator of the patient 5-year survival rate in the R package survival^98^ (**Figure 4C**). Multivariate survival analysis was performed to show factors significantly (*p* < 0.05) correlated with patient survival with 95% confidence interval (**Figure 4D**).

### Feedforward loop analysis

We built a network with three types of nodes: motifs, TET1/DNMT3A, coding genes. We have defined promoters as the region −1000bp and +500bp from transcription start sites (TSS) of protein coding genes (including TET1 and DNMT3A) from Gencode v19^93^, as previously described. A directed edge is defined if the source node has an occurrence sites at the promoter of the target nodes. For TET1 and DNMT3A, occurrence site is defined by ChIP-seq data previously measured in hESC and NCCIT cells, respectively. For motifs, the occurrence site is defined by FIMO with *p* < 10^−5^. When coding gene is a target, we first check if the gene is a known transcription factor, then define its binding site by FIMO with *p* < 10^−5^. Finally, tracks are visualized in integrated genome viewer and the methylation track is provided by WGBS of H1 from The Epigenomics Roadmap Project^14^.

## Supporting information

Supplementary Figures (high resolution)

## Author contributions

M.W. performed the computational analyses and interpreted the results. K.Z., V.N. and J.W.W. helped discovered the *de novo* motifs. K.Z. designed the motif mering and contributed to finding motif occurrence. S.F., Y.C. and L.Z. contributed to TCGA methylation analysis. V.N. and J.W. contributed to SNP analysis. C.L. Z.C. contributed to functional annotation of motifs. W.W. conceived the study, supervised the analyses and interpreted the results. M.W. and W.W. wrote the paper with contribution from all authors.

## Acknowledgement

This project is partially supported by NIH (U54HG006997 and R01HG009626) and CIRM (RB5 07012).

## Supplementary Figures

**Figure S1.**
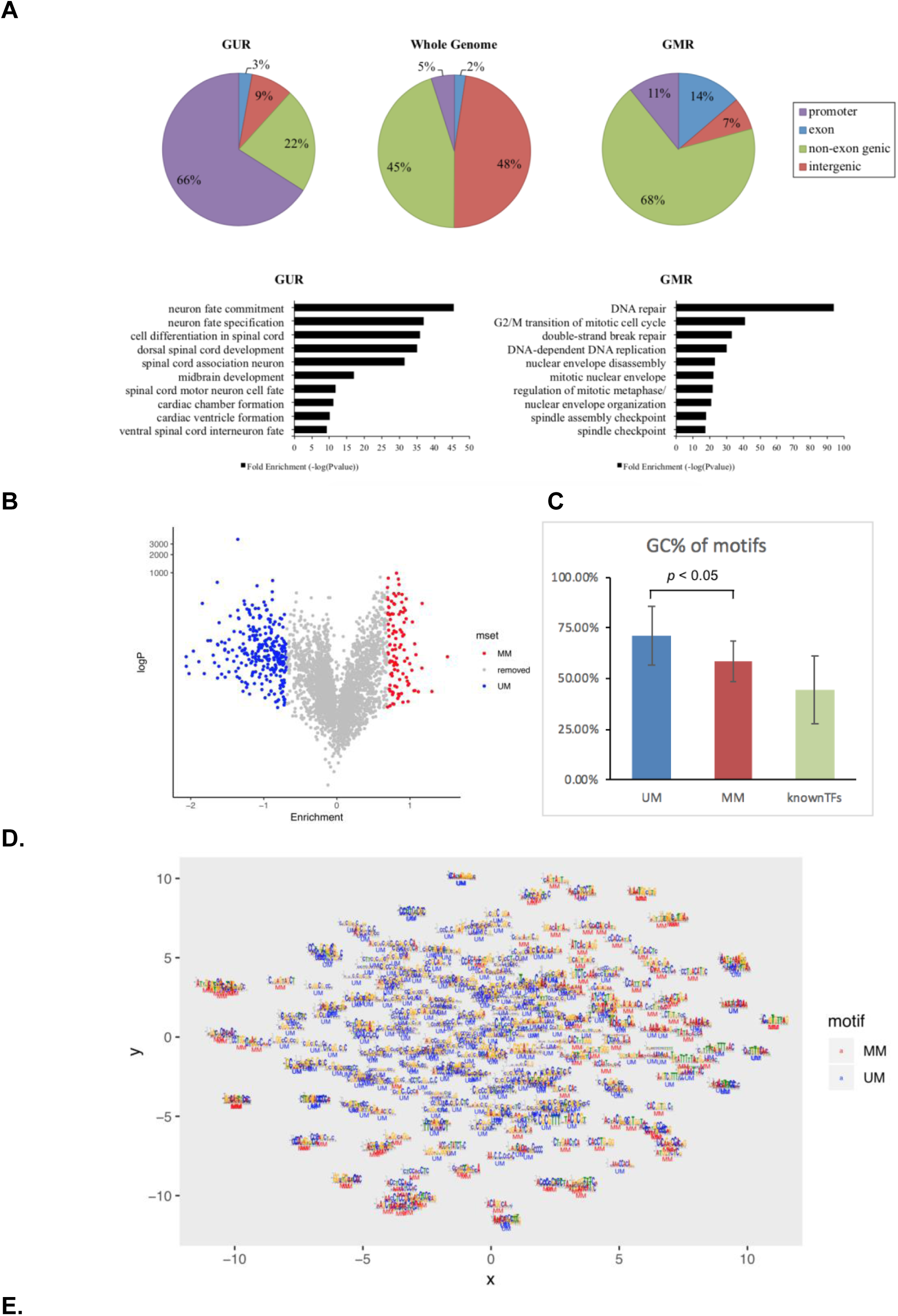
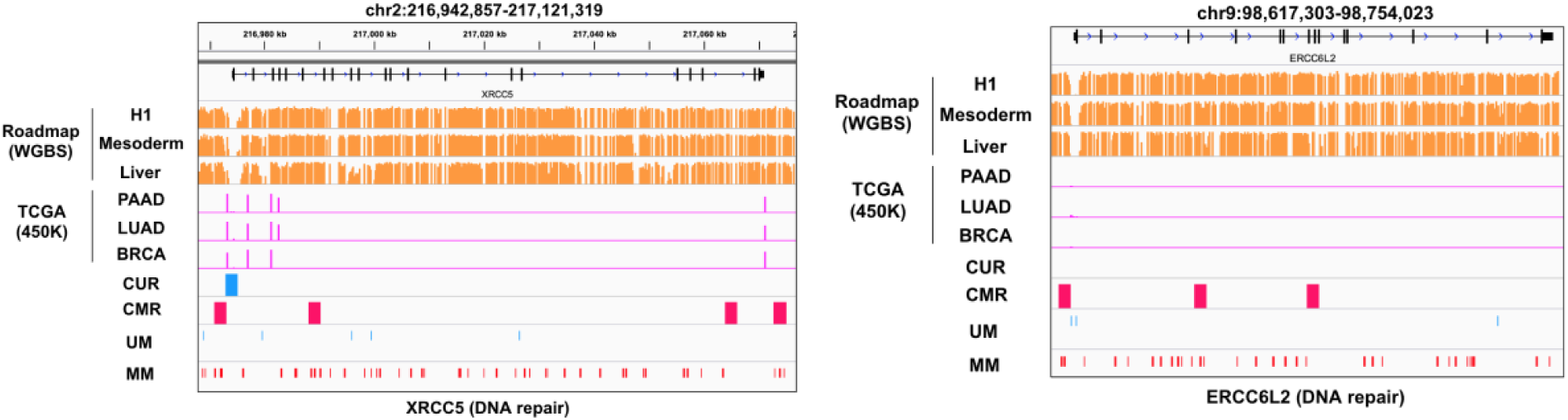
Characterization of the identified motifs and regions. **A**. Gene ontology analysis and genomic location of CUR and CMR compared against the whole genome. **B.** Volcano plot of 313 top cluster filtered by logP>10 and fold-change enrichment>2. **C**. Motif GC content of top 313 unmethylation motifs (UM), methylation motifs (MM) and known motifs curated from hocomoco^29^. **D**. tSNE^99^ plot showing sequence similarity among 313 motifs, pairwise distance calculation described in method sequence alignment. **E.** Methylation correlated with CUR, CMR and motif locations at XRCC5 and ERCC6L2, two genes associated with DNA repair.

**Figure S2.**
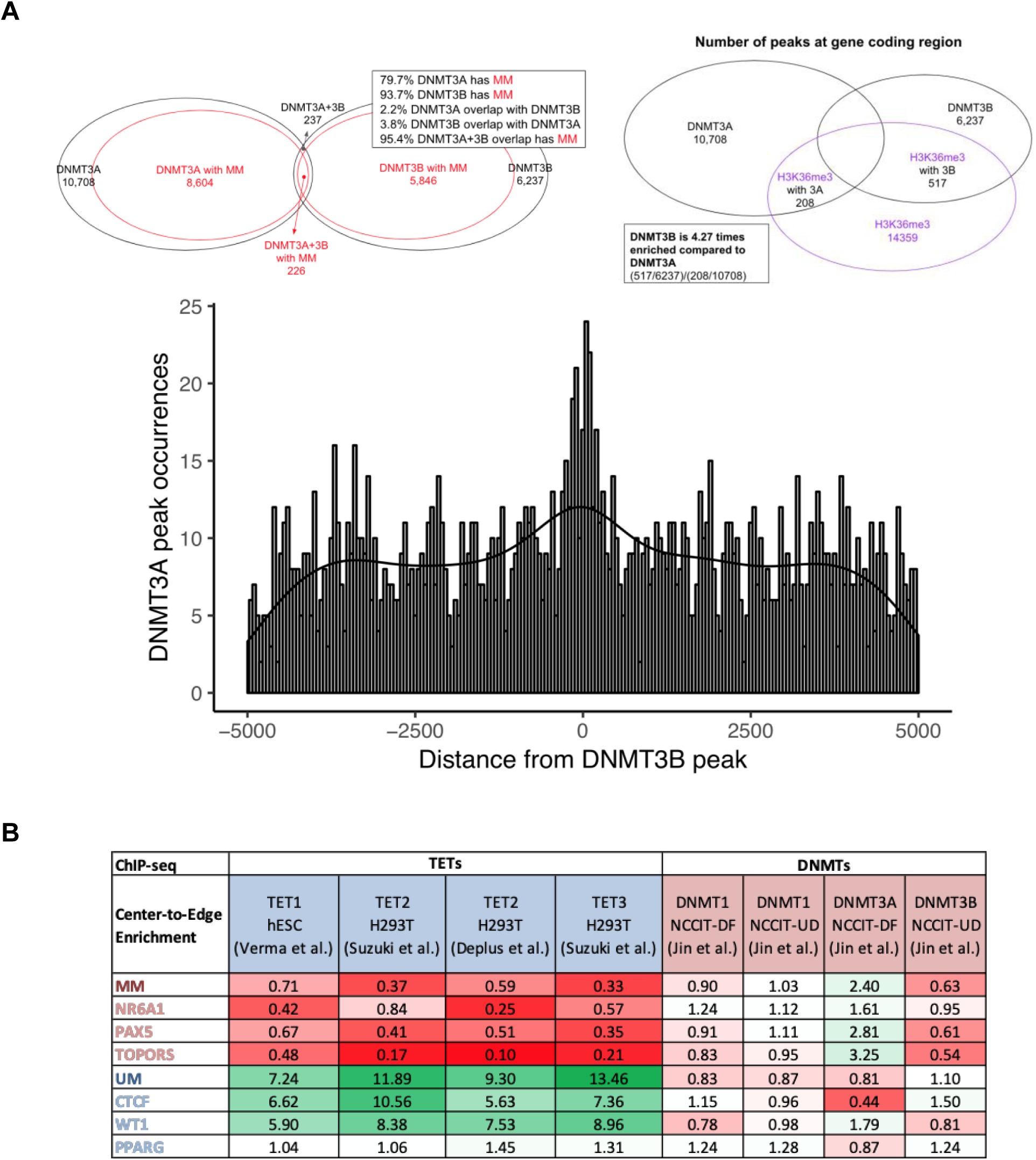
Identified motifs marks methylation level **A.** methylation motif co-occur with DNMT1 and DNMT3B ChIP-seq^45^ peaks in differentiated NCCIT cell, while unmethylation motif co-occur with TET ChIP-seq^46^ peaks in human embryonic stem cells. Lower panel shows the histogram of DNMT3A peak counts 5000bp nearby DNMT3B peaks, with 50bp bin width. **B.** Details on center-to-edge enrichment of motifs and known TFs in respect to TETs and DNMTs ChIP-seq peaks.

**Figure S3.**
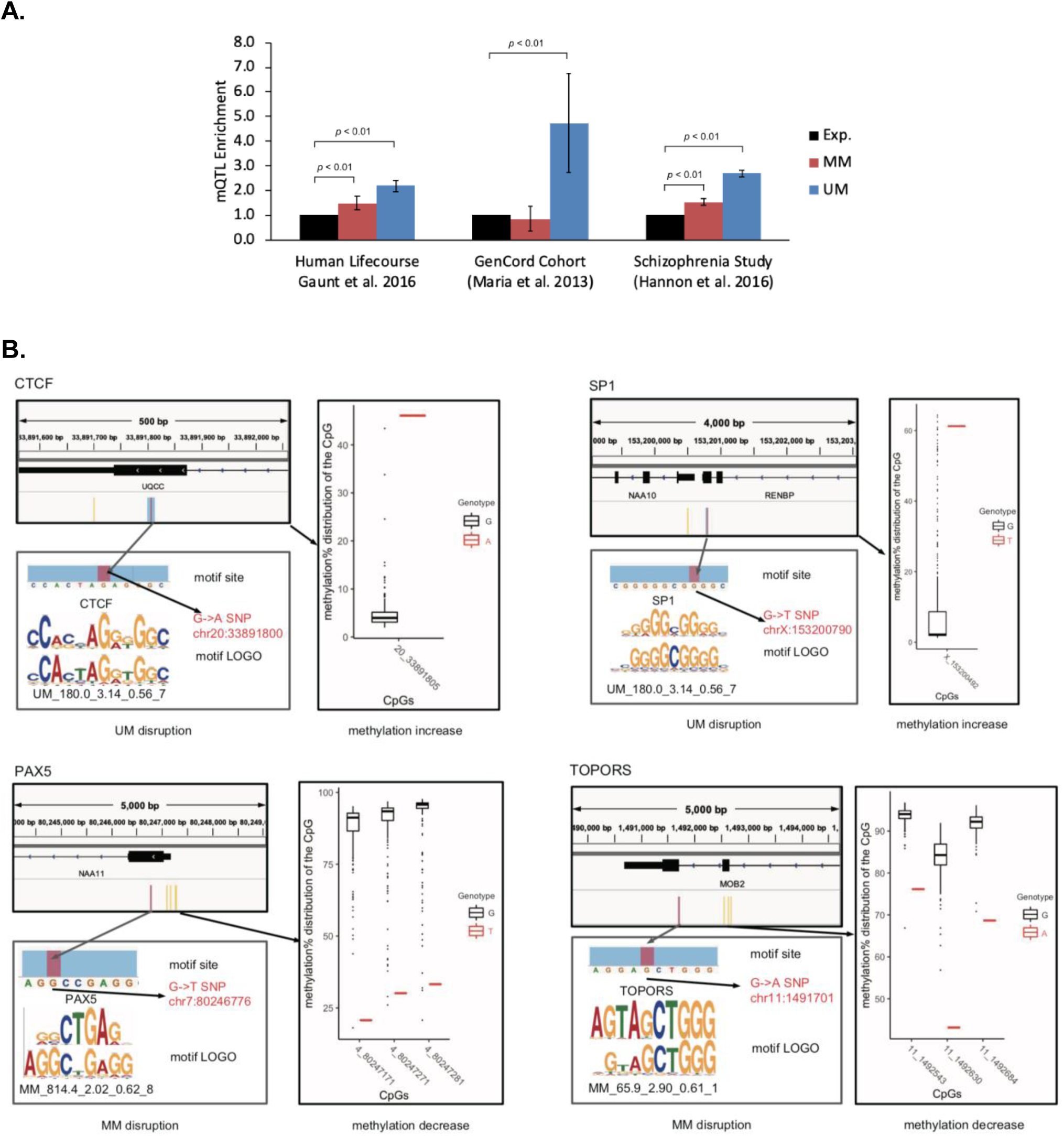
Identified motif occurrences overlaps with TCGA functional SNPs. **A.** Enrichment of mQTL at UM and MM occurrence site using three additional studies, namely human life course study^63^, GenCord Cohort study^64^ and a Schizophrenia study^65^. Enrichment of mQTL are defined as observed mQTL ratio over expected ratio, with error bars showing standard deviation across samples (Left: across five stages in human life course; Middle: across three cell types; Right: across three adult human brain regions) and *p* < 0.01 t-test are marked (See **method** for details). **B.** More examples UMs and MMs matched to known motifs CTCF, SP1, PAX5 and TOPORS disrupted by somatic mutations show correlation with local methylation alteration.

**Figure S4.**
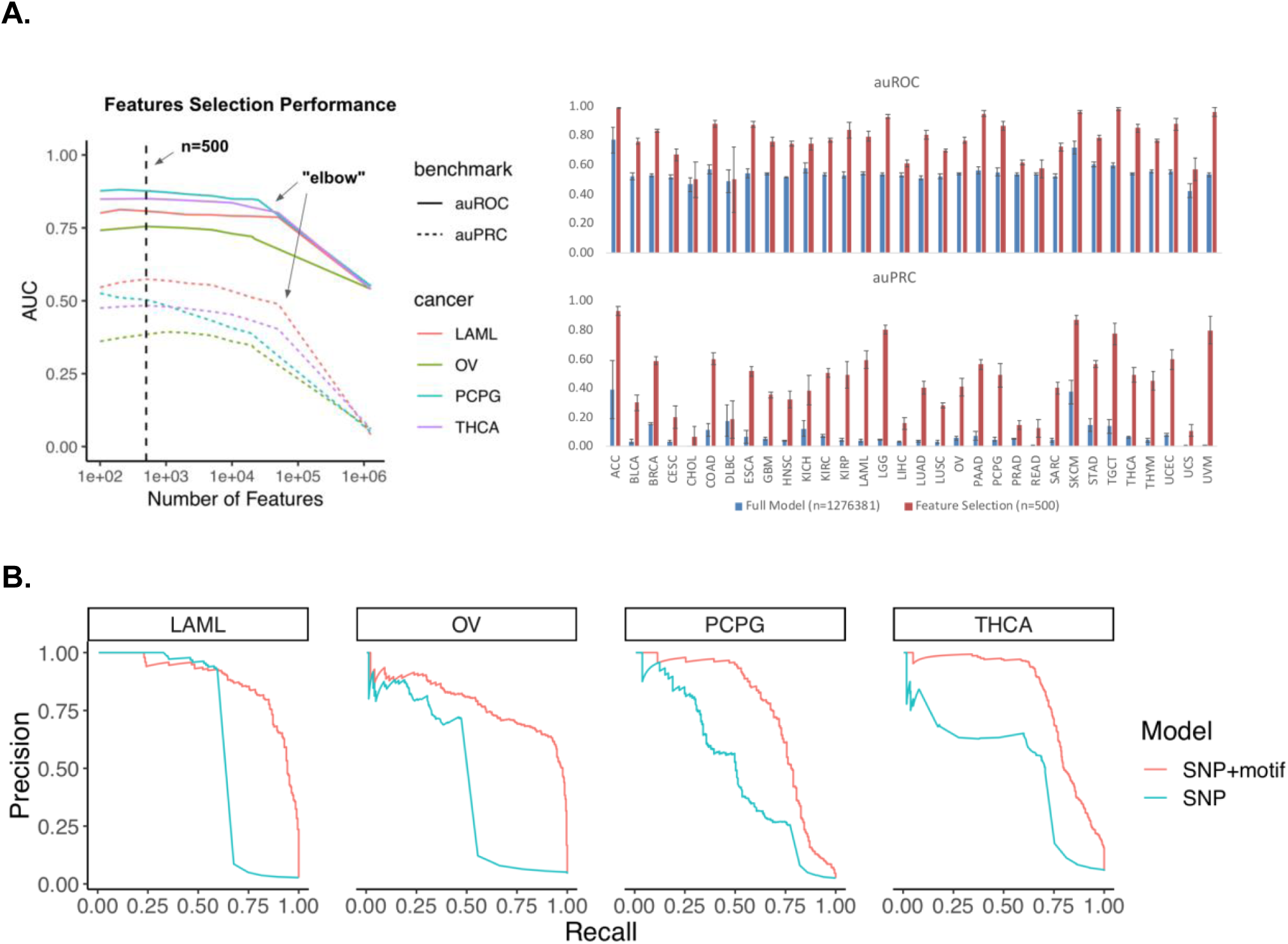
Motifs disrupted by SNPs predicts cancer subtype and survival **A**. Feature selection reduces feature number while improve performance for SNP-only diagnosis model. Elbow indicates the number of features having positive feature importance scores. **B**. Performance evaluation of models predicting patient cancer subtype using auROC metric.

