## Supplementary Figures (high resolution) for "Identification of DNA motifs that regulate DNA methylation"

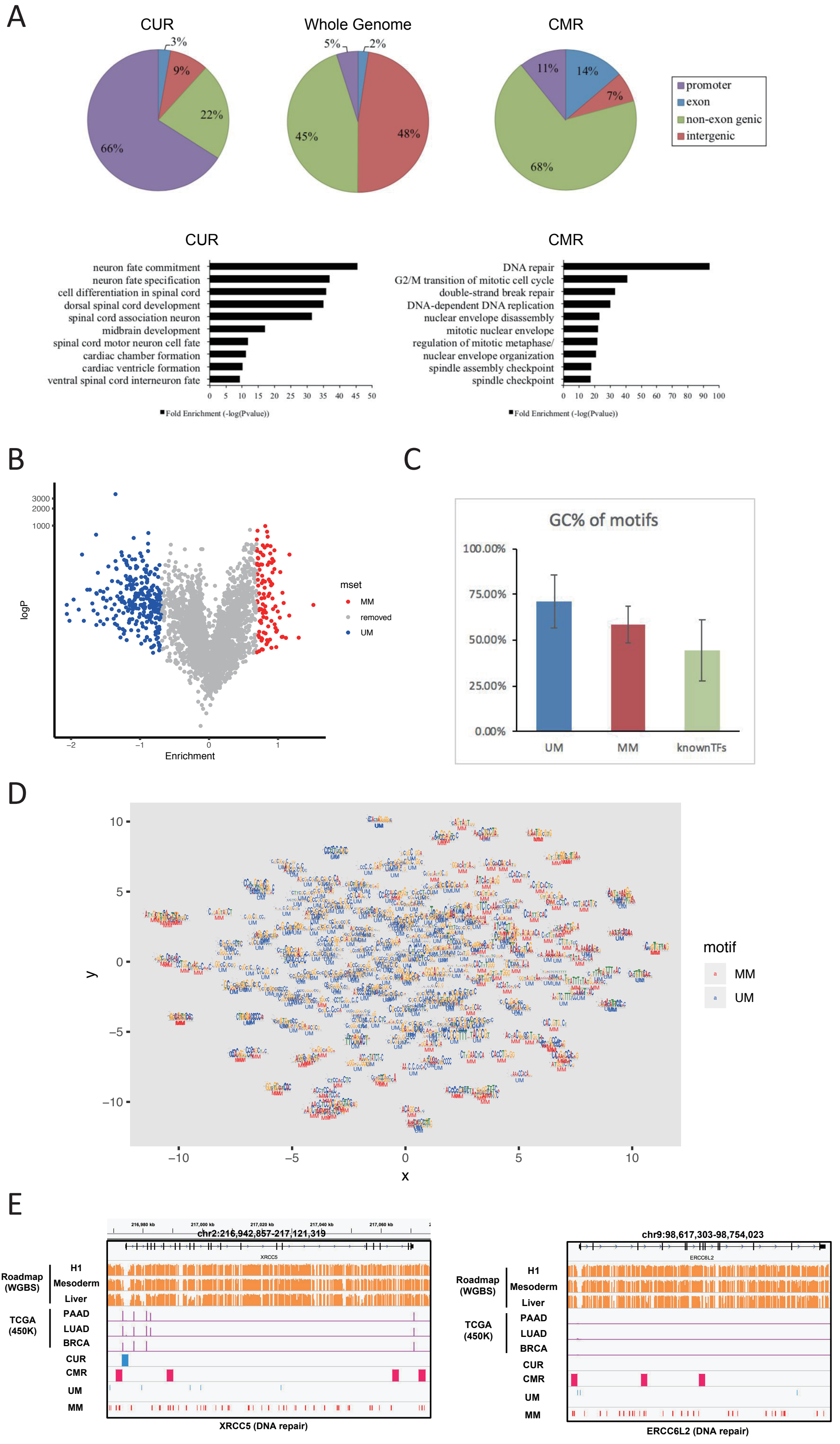

A
CUR
3%
9%
22%
66%
Whole Genome
5%
2%
45%
48%
CMR
11%
14%
7%
68%
promoter
exon
non-exon genic
intergenic
CUR
neuron fate commitment
neuron fate specification
cell differentiation in spinal cord
dorsal spinal cord development
spinal cord association neuron
midbrain development
spinal cord motor neuron cell fate
cardiac chamber formation
cardiac ventricle formation
ventral spinal cord interneuron fate
0 5 10 15 20 25 30 35 40 45 50
Fold Enrichment (-log(Pvalue))
CMR
DNA repair
G2/M transition of mitotic cell cycle
double-strand break repair
DNA-dependent DNA replication
nuclear envelope disassembly
mitotic nuclear envelope
regulation of mitotic metaphase/
nuclear envelope organization
spindle assembly checkpoint
spindle checkpoint
0 10 20 30 40 50 60 70 80 90 100
Fold Enrichment (-log(Pvalue))
B
3000
2000
1000
logP
−2 −1 0 1
Enrichment
mset
MM
removed
UM
C
GC% of motifs
100.00%
75.00%
50.00%
25.00%
0.00%
UM
MM
knownTFs
D
10
5
0
−5
−10
y
−10 −5 0 5 10
x
motif
MM
UM
E
216,980 kb
217,000 kb
217,020 kb
217,040 kb
217,060 kb
chr2:216,942,857-217,121,319
XRCC5
Roadmap (WGBS)
H1
Mesoderm
Liver
TCGA (450K)
PAAD
LUAD
BRCA
CUR
CMR
UM
MM
XRCC5 (DNA repair)
chr9:98,617,303-98,754,023
ERCC6L2
Roadmap (WGBS)
H1
Mesoderm
Liver
TCGA (450K)
PAAD
LUAD
BRCA
CUR
CMR
UM
MM
ERCC6L2 (DNA repair)

A

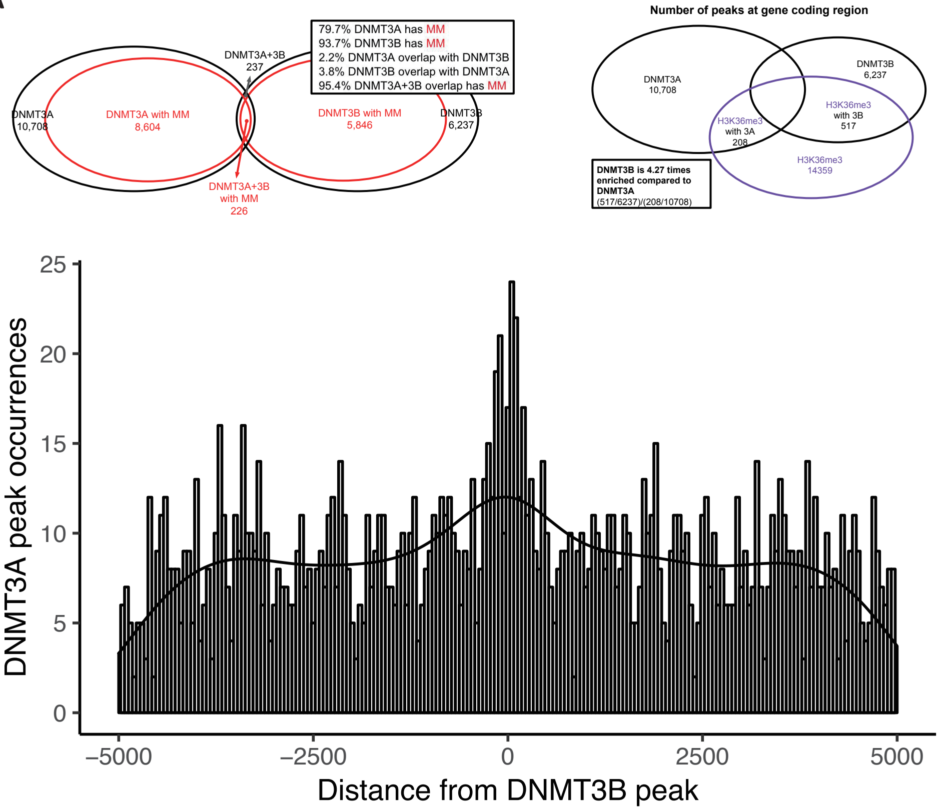


B

| ChIP-seq | TETs | | | | DNMTs | | | |
|---|---|---|---|---|---|---|---|---|
| Center-to-Edge Enrichment | TET1 hESC (Verma et al.) | TET2 H293T (Suzuki et al.) | TET2 H293T (Deplus et al.) | TET3 H293T (Suzuki et al.) | DNMT1 NCCIT-DF (Jin et al.) | DNMT1 NCCIT-UD (Jin et al.) | DNMT3A NCCIT-DF (Jin et al.) | DNMT3B NCCIT-UD (Jin et al.) |
| MM | 0.71 | 0.37 | 0.59 | 0.33 | 0.90 | 1.03 | 2.40 | 0.63 |
| NR6A1 | 0.42 | 0.84 | 0.25 | 0.57 | 1.24 | 1.12 | 1.61 | 0.95 |
| PAX5 | 0.67 | 0.41 | 0.51 | 0.35 | 0.91 | 1.11 | 2.81 | 0.61 |
| TOPORS | 0.48 | 0.17 | 0.10 | 0.21 | 0.83 | 0.95 | 3.25 | 0.54 |
| UM | 7.24 | 11.89 | 9.30 | 13.46 | 0.83 | 0.87 | 0.81 | 1.10 |
| CTCF | 6.62 | 10.56 | 5.63 | 7.36 | 1.15 | 0.96 | 0.44 | 1.50 |
| WT1 | 5.90 | 8.38 | 7.53 | 8.96 | 0.78 | 0.98 | 1.79 | 0.81 |
| PPARG | 1.04 | 1.06 | 1.45 | 1.31 | 1.24 | 1.28 | 0.87 | 1.24 |

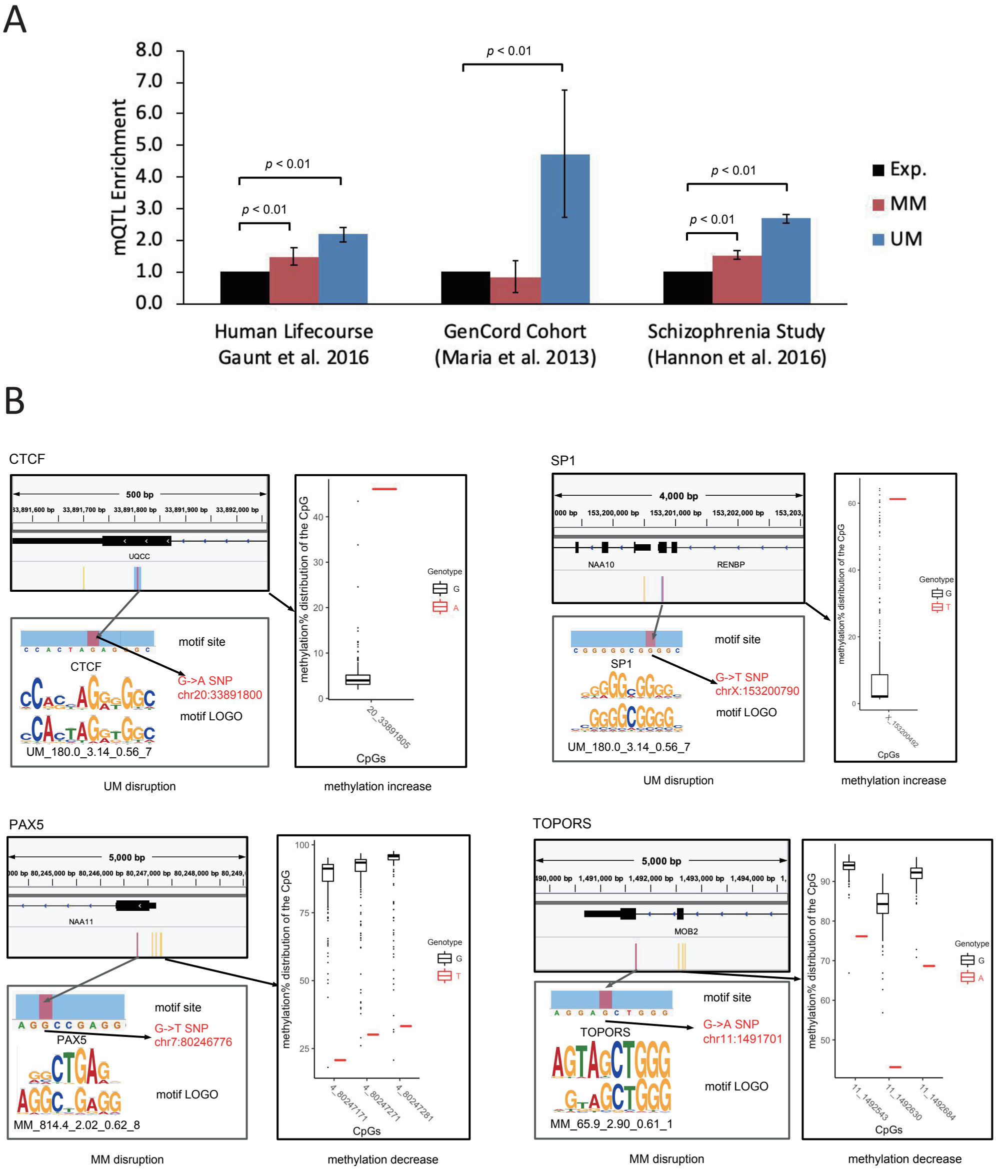
A
mQTL Enrichment
8.0
7.0
6.0
5.0
4.0
3.0
2.0
1.0
0.0
p < 0.01
p < 0.01
p < 0.01
p < 0.01
p < 0.01
Exp.
MM
UM
Human Lifecourse
Gaunt et al. 2016
GenCord Cohort
(Maria et al. 2013)
Schizophrenia Study
(Hannon et al. 2016)
B
CTCF
500 bp
UQCC
motif site
CTCF
G->A SNP
chr20:33891800
motif LOGO
UM_180.0_3.14_0.56_7
UM disruption
methylation% distribution of the CpG
Genotype
G
A
20_33891805
CpGs
methylation increase
SP1
4,000 bp
NAA10
RENBP
motif site
SP1
G->T SNP
chrX:153200790
motif LOGO
UM_180.0_3.14_0.56_7
UM disruption
Genotype
G
T
X_153200492
CpGs
methylation increase
PAX5
5,000 bp
NAA11
motif site
PAX5
G->T SNP
chr7:80246776
motif LOGO
MM_814.4_2.02_0.62_8
MM disruption
Genotype
G
T
4_80247171
4_80247271
4_80247281
CpGs
methylation decrease
TOPORS
5,000 bp
MOB2
motif site
TOPORS
G->A SNP
chr11:1491701
motif LOGO
MM_65.9_2.90_0.61_1
MM disruption
Genotype
G
A
11_1492543
11_1492630
11_1492684
CpGs
methylation decrease

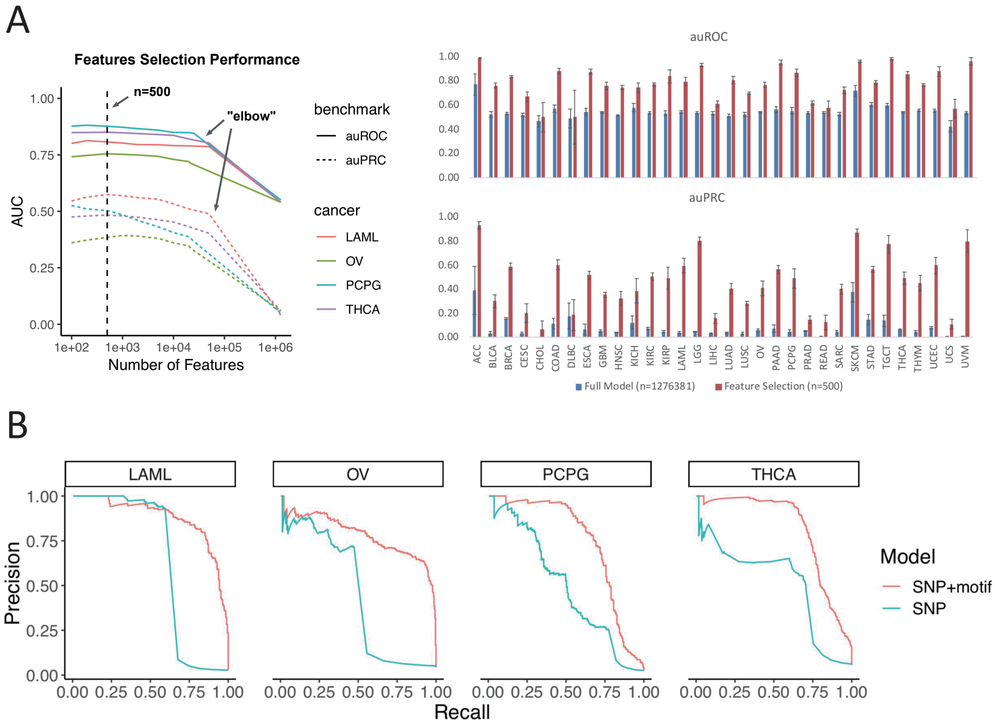
A
Features Selection Performance
n=500
"elbow"
AUC
Number of Features
1.00
0.75
0.50
0.25
0.00
1e+02
1e+03
1e+04
1e+05
1e+06
benchmark
auROC
auPRC
cancer
LAML
OV
PCPG
THCA
auROC
auPRC
ACC BLCA BRCA CESC CHOL COAD DLBC ESCA GBM HNSC KICH KIRC KIRP LAML LGG LIHC LUAD LUSC OV PAAD PCPG PRAD READ SARC SKCM STAD TGCT THCA THYM UCEC UCS UVM
Full Model (n=1276381)
Feature Selection (n=500)
B
LAML
OV
PCPG
THCA
Precision
Recall
0.00 0.25 0.50 0.75 1.00
Model
SNP+motif
SNP
